# MTAG: Multi-Trait Analysis of GWAS

**DOI:** 10.1101/118810

**Authors:** Patrick Turley, Raymond K. Walters, Omeed Maghzian, Aysu Okbay, James J. Lee, Mark Alan Fontana, Tuan Anh Nguyen-Viet, Robbee Wedow, Meghan Zacher, Nicholas A. Furlotte, 23andMe Research Team, Social Science Genetic Association Consortium, Patrik Magnusson, Sven Oskarsson, Magnus Johannesson, Peter M. Visscher, David Laibson, David Cesarini, Benjamin M. Neale, Daniel J. Benjamin

**Affiliations:** Broad Institute, 415 Main Street, Cambridge, Massachusetts 02142, United States; Analytic and Translational Genetics Unit, Massachusetts General Hospital, 185 Cambridge Street, Cambridge, Massachusetts 02114, United States; Department of Economics, Harvard University, 1805 Cambridge Street, Cambridge, Massachusetts 02138, United States; Department of Complex Trait Genetics, Vrije Universiteit Amsterdam, De Boelelaan 1085, Amsterdam, 1081 HV, Netherlands; Department of Psychology, University of Minnesota, 75 E River Road, Minneapolis, Minnesota 55455, United States; Hospital for Special Surgery, 535 E 70th Street, New York, New York 10021, United States; Center for Economic and Social Research, University of Southern California, 635 Downey Way, Los Angeles, California 90089, United States.; Institute for Behavioral Genetics, University of Colorado Boulder, 1480 30th Street, Boulder, 80309, Colorado, United States.; Institute of Behavioral Science, University of Colorado Boulder, 1440 15th Street, Boulder, 80309, Colorado, United States.; Department of Sociology, University of Colorado Boulder, UCB 327, Boulder, 80309, Colorado, United States.; Department of Sociology, Harvard University, 33 Kirkland Street, Cambridge, Massachusetts, 02138, United States.; 23andMe, Inc., Mountain View, California 94043, United States; Institutionen för Medicinsk Epidemiologi och Biostatistik, Karolinska Institutet, 171 77 Stockholm, Sweden; Department of Government, Uppsala Universitet, 751 20 Uppsala, Sweden; Department of Economics, Stockholm School of Economics, 113 83 Stockholm, Sweden; Institute for Molecular Bioscience, The University of Queensland, QLD 4072, Brisbane, Queensland, Australia; Queensland Brain Institute, The University of Queensland, QLD 4072, Brisbane, Queensland, Australia; National Bureau of Economic Research, 1050 Massachusetts Avenue, Cambridge, Massachusetts 02138, United States; Department of Economics and Center for Experimental Social Science, New York University, 19 W 4th Street, New York, New York 10012, United States; Institutet för Näringslivsforskning, 114 53 Stockholm, Sweden; Department of Economics, University of Southern California, 3620 South Vermont Avenue, Kaprielian Hall, 300 Los Angeles, CA 90089, United States

## Abstract

We introduce Multi-Trait Analysis of GWAS (MTAG), a method for joint analysis of summary statistics from GWASs of different traits, possibly from overlapping samples. We apply MTAG to summary statistics for depressive symptoms (*N_eff_* = 354,862), neuroticism (*N* = 168,105), and subjective well-being (*N* = 388,538). Compared to 32, 9, and 13 genome-wide significant loci in the single-trait GWASs (most of which are themselves novel), MTAG increases the number of loci to 64, 37, and 49, respectively. Moreover, association statistics from MTAG yield more informative bioinformatics analyses and increase variance explained by polygenic scores by approximately 25%, matching theoretical expectations.

## 1 INTRODUCTION

The standard approach in genetic-association studies is to analyze a single trait. Such studies do not exploit information contained in summary statistics from genome-wide association studies (GWASs) of related traits. In this paper, we develop a method, Multi-Trait Analysis of GWAS (MTAG), which enables joint analysis of multiple traits, thus boosting statistical power to detect genetic associations for *each* trait.

Compared to the many existing multi-trait methods,^1–5^ MTAG has a unique combination of four features that make it potentially useful in many settings. First, it can be applied to GWAS summary statistics (without access to individual-level data) from an arbitrary number of traits. Second, the summary statistics need not come from independent discovery samples: MTAG uses bivariate linkage disequilibrium (LD) score regression^6^ to account for (possibly unknown) sample overlap between the GWAS results for different traits. Third, MTAG generates trait-specific effect estimates for each single-nucleotide polymorphism (SNP). Finally, even when applied to many traits, MTAG is computationally quick because every step has a closed-form solution.

The MTAG estimator is a generalization of inverse-variance-weighted meta-analysis that takes summary statistics from single-trait GWASs and outputs trait-specific association statistics. The resulting *P* values can be interpreted and used like *P* values from a single-trait GWAS, e.g., to prioritize SNPs for subsequent analyses such as biological annotation or to construct polygenic scores.

The key assumption of MTAG is that all SNPs share the same variance-covariance matrix of effect sizes across traits. This assumption is strong and is violated in many circumstances, most intuitively in scenarios where some SNPs influence only a subset of the traits. Even if this assumption is not satisfied, however, we show analytically that MTAG is a consistent estimator and that its effect estimates *always* have a lower genome-wide mean squared error than the corresponding single-trait GWAS estimates. Hence, polygenic scores constructed from MTAG results are expected to outperform GWAS-based predictors very generally.

The main potential problem arises for SNPs that are truly null for one trait but non-null for another trait. For such SNPs, MTAG’s effect-size estimates for the first trait are biased away from zero, leading to an increased rate of false positives. We derive an analytic formula for the resulting false discovery rate (FDR), given any specified mixture-normal distribution of effect sizes (including multivariate spike-and-slab distributions), and we illustrate how the formula can be used to probe the credibility of MTAG-identified loci.

To demonstrate the utility of MTAG empirically, we analyze three traits: depressive symptoms (DEP, *N_eff_* = 354,862), neuroticism (NEUR, *N* = 168,105), and subjective well-being (SWB, *N* = 388,538). Prior GWASs of each of these traits have identified only a handful of loci.^7–11^ Because of the high genetic correlations between the three traits—in our data, roughly 0.7 in absolute value between each pair—some papers have conducted cross-trait analyses to replicate findings for one of the traits^11^ or joint meta-analysis to identify new loci.^5^ We apply MTAG to these traits because we expected the gains in power would be substantial, violations of MTAG’s assumptions would be limited, and the substantive results would be of interest.

Finally, we compare MTAG to the three existing multi-trait methods we are aware of that can be applied to GWAS summary statistics from an arbitrary number of traits with unknown sample overlap.^12,13^ We find that MTAG has greater power across a wide range of simulation scenarios and in two separate applications to real data.

## RESULTS

### Overview of MTAG

The key idea underlying MTAG is that when GWAS estimates from different traits are correlated, the effect estimates for each trait can be improved by appropriately incorporating information contained in the GWAS estimates for the *other* traits.

Correlation between GWAS estimates can arise for two reasons. First, the traits may be genetically correlated, in which case the *true effects* of the SNPs are correlated across traits. Second, the *estimation error* of the SNPs’ effects may be correlated across traits. Such correlation will occur if (a) the phenotypic correlations are non-zero and there is sample overlap across traits, or if (b) biases in the SNP-effect estimates (e.g., population stratification or cryptic relatedness) have correlated effects across traits. MTAG boosts statistical power by incorporating information about these two sources of correlation.

### MTAG Framework

In the framework that follows, all traits and genotypes are standardized to have mean zero and variance one. For SNP *j*, we denote the vector of marginal (i.e., not controlling for other SNPs), true effects on each of the *T* traits by ***β**_j_*. We treat these true effects as random effects with E(***β**_j_*) = 0 and Var(***β**_j_*) = **Ω**. If the true effects are correlated across traits, then the off-diagonal elements of **Ω** are non-zero. MTAG’s key assumption is that **Ω** is homogeneous across SNPs, i.e., it does not depend on *j*.

We denote the vector of GWAS estimates of SNP *j*’s effects on the traits by 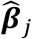. We assume that the GWAS estimates are unbiased, 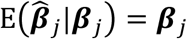, and we denote the variance-covariance matrix of their estimation error by 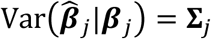. The off-diagonal elements of **Σ**_*j*_ are non-zero whenever the estimation errors are correlated.

MTAG is the efficient generalized method of moments (GMM) estimator based on the moment condition

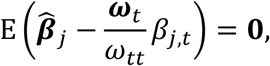

where ***ω**_j_* is a vector equal to the *t*^th^ column of **Ω** and *ω_tt_* is a scalar equal to the *t*^th^ diagonal element of **Ω**. This equality is a necessary condition derived from the best linear prediction of the vector of GWAS estimates, 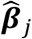, from the SNP’s true effect on a single trait, *β_j,t_*.

The MTAG estimator is a weighted sum of the GWAS estimates:

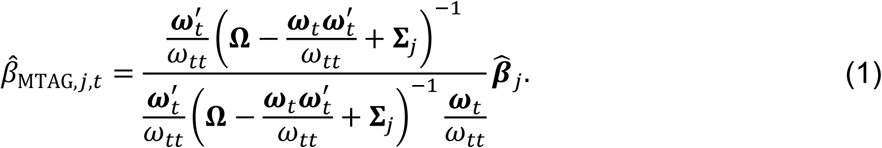

It is a consistent and asymptotically normal estimator for *β_j,t_* (**Supplemental Note**).

There are several useful special cases of MTAG (see **Online Methods**). When all estimates are for the same trait (implying 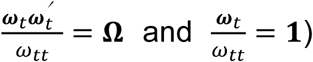, equation (1) simplifies to: 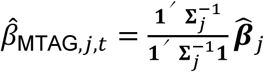. When the GWAS estimates are obtained from non-overlapping samples (i.e., **Σ**_*j*_ is diagonal), this formula specializes to the well-known formula for inverse-variance-weighted meta-analysis. When the genetic correlations across all traits are zero and there is no sample overlap (i.e., both Ω and **Σ**_*j*_ are diagonal), the MTAG estimates are identical to the GWAS estimates. This equivalence is intuitive, since it corresponds exactly to the case of no correlation between the GWAS estimates that can be leveraged.

To make equation (1) operational, we use consistent estimates of **Σ**_*j*_ and **Ω**, described next (see **Supplementary Note** for details).

#### Estimation of Σ_j_

In standard meta-analysis, the diagonal elements of 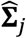 would be constructed using the squared standard errors from the GWAS results, and the off-diagonal elements of 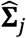 would be set to zero. In MTAG, however, we want to allow the estimation error to include bias (in addition to sampling variation) and to be correlated across the GWAS estimates.

Therefore, MTAG proceeds by running linkage disequilibrium (LD) score regressions^14^ on the GWAS results and using the estimated intercepts to construct the diagonal elements of 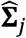. Next, bivariate LD score regressions^6^ are run using each pair of GWAS results, and the estimated intercepts are used to construct the off-diagonal elements of 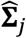. Under the assumptions of LD score regression (including that the LD reference sample and GWAS samples are all drawn from the same population), the resulting matrix 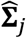 captures all relevant sources of estimation error, including not only sampling variation but also population stratification, unknown sample overlap, and cryptic relatedness. Because the LD-score-intercept adjustment is already built into the MTAG estimates, MTAG-generated association results do not require further adjustment for these biases.

#### Estimation of Ω

We estimate 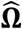 by method of moments using the moment condition

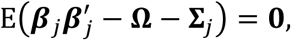

with 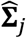 substituted in place of **Σ**_*j*_. This is the step that relies on the homogeneous-**Ω** assumption: the assumption justifies using all SNPs when estimating 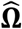.

#### Summary

The MTAG results for SNP *j* are obtained in three steps: (i) estimate the variance-covariance matrix of the GWAS estimation error, 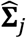, by using a sequence of LD score regressions, (ii) estimate the variance-covariance matrix of the SNP effects, 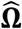, using method of moments, and (iii) for each SNP, substitute these estimates into equation (1). We have made available for download a Python command line tool that runs our MTAG estimation procedure (see URLs). Because each of the above steps has a closed-form solution, genome-wide analyses using the MTAG software run quickly (see **Online Methods**).

### Theoretical Analysis of MTAG’s Performance

This section briefly discusses three analytic formulas we have derived regarding the expected performance of MTAG for each trait: its mean squared error (MSE) across SNPs, its statistical power to detect a true single-SNP association, and its false discovery rate (FDR) (**Online Methods**). All the formulas hold for an arbitrary number of traits. **Supplementary Note** contains illustrative calculations. The formulas depend on **Ω** and **Σ**_*j*_ but can be approximated in applications using 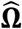 and 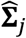.

The MSE formula is very general: it holds under any distribution of effect sizes, including distributions that violate the homogeneous-**Ω** assumption. The formula implies that for each trait, the MTAG estimates *always* have a lower genome-wide MSE than corresponding GWAS estimates. That in turn suggests that polygenic predictors constructed from MTAG results are likely to outperform GWAS-based predictors very generally.

The power and FDR formulas (in contrast to the fully general MSE formula) assume that the true effect sizes ***β**_j_* are drawn from some known mean-zero mixture of multivariate normal distributions. This class of distributions includes multivariate spike-and-slab distributions and other fat-tailed distributions that may be relevant in applications of MTAG.

### Potential Biases in MTAG’s Test Statistics

The derivation of MTAG relies on three important assumptions: (1) **Ω** is homogeneous across SNPs, (2) sampling variation in 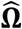 and 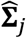 can be ignored, and (3) the off-diagonal elements of 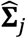 (estimated by bivariate LD score regression) accurately capture sample overlap. In light of each assumption, here we probe when and to what extent MTAG’s test statistics for individual-SNP associations may be biased.

#### Homogeneous-Ω assumption

If the homogeneous-**Ω** assumption is violated, then there are different types of SNPs with different **Ω**’s. Because MTAG combines the GWAS estimates using the genome-wide (i.e., across-SNP) variance-covariance matrix, in general the MTAG estimates will be biased in finite samples. For a type of SNP that is null for one trait but non-null for other traits, the effect estimate on the first trait will be biased away from zero. For that reason, the FDR will be inflated.

Replication is the best way to assess the credibility of individual-SNP associations. In addition, their credibility can be probed using the FDR formula, computed under plausible assumptions about genetic architecture. In our application below, we calculate what we call *maxFDR*, which is an upper bound for the FDR under certain assumptions (**Online Methods**). In particular, we assume that the effect-size distribution is a multivariate spike-and-slab distribution in which at least 10% of SNPs are non-null for each trait. Illustrative calculations indicate that a trait’s maxFDR can become high when the GWAS for the trait is low powered while the GWAS for another trait is higher powered (**Supplementary Note**).

#### Sampling variation in 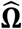 and 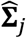 ignored

To assess the magnitude of the finite-sample bias in MTAG’s standard errors from ignoring sampling variation in 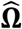 and 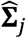, we simulate GWAS summary statistics for up to *T* = 20 traits and apply MTAG using 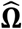 and 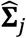 (as in any real-data application of MTAG). We then calculate the inflation of the mean *χ*^2^-statistic, defined relative to what the mean *χ*^2^-statistic would be if the true values **Ω** and **Σ**_*j*_ were used. Figures 1a and 1b plots the inflation as a function of *T*, where each GWAS has mean *χ*^2^-statistic of 1.1, 1.4, or 2.0. The effect-size correlation between every pair of traits is *r_β_* = 0 (Figure 1a) or *r_β_* = 0.7 (Figure 1b); we set the correlation in estimation error between every pair of traits to *r_ε_* = 0 in these simulations. The figure shows that inflation increases roughly linearly in the number of traits. The bias is larger when the GWASs are lower powered and when *r_β_* is smaller. Our application to DEP, NEUR, and SWB (discussed below) corresponds roughly to a mean *χ*^2^-statistic of 1.4 with *T* = 3 in Figure 1b. In that setting, inflation is negligible. Even when inflation is largest—the low-powered GWAS with *T* = 20 in Figure 1a—it is only 3%.

**Fig. 1.**
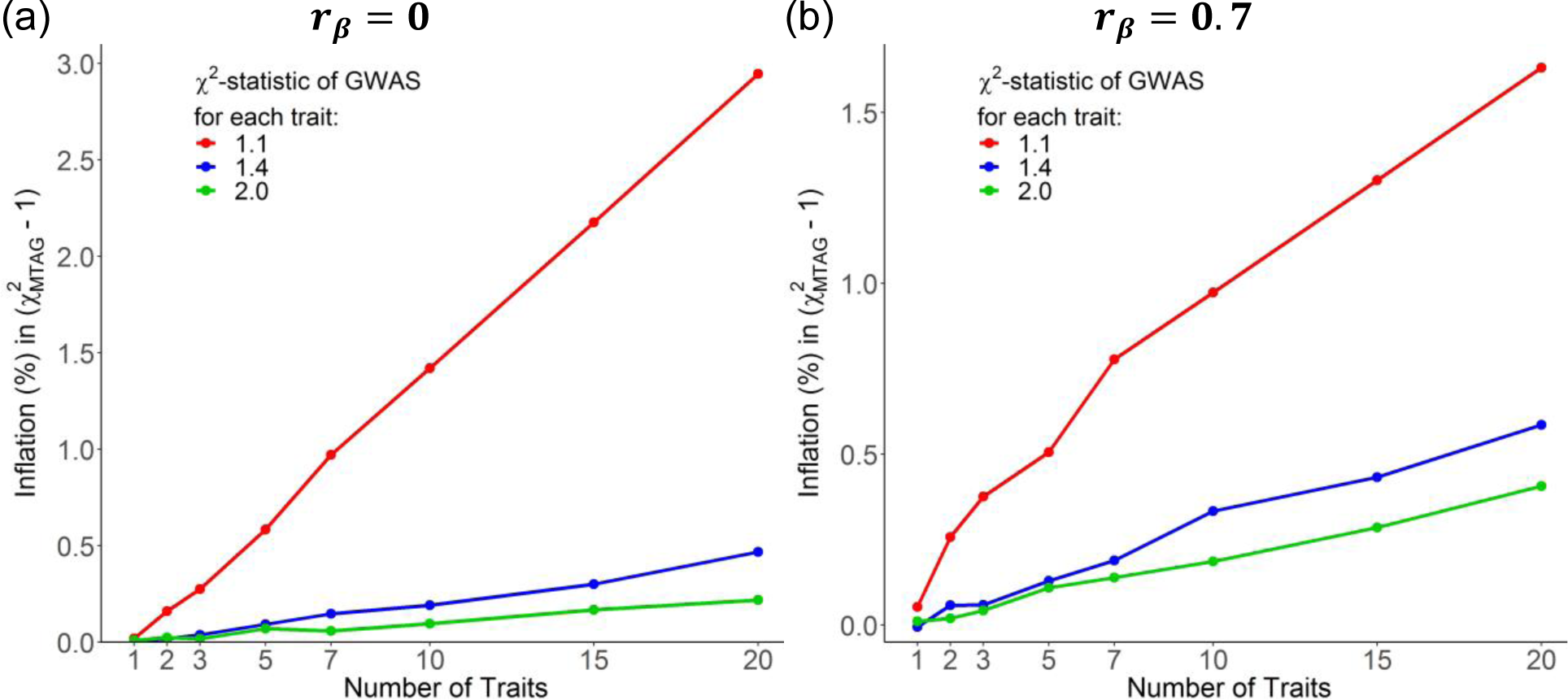
Bias in standard errors from ignoring sampling variation in 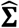 and 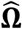. The *y*-axis is the percent increase in (*χ*^2^ − 1) of the MTAG test statistics from using estimated values of **Σ** and **Ω** rather than the true values. Each line corresponds to results from applying MTAG to identically powered single-trait GWASs of *T* traits. For every pair of traits, the correlation in true effect sizes is (**a**) *r_β_* = 0, (**b**) *r_β_* = 0.7. Complete results for the full set of simulation scenarios can be found in **Supplementary Note**.

These simulations suggest that in most realistic applications of MTAG, the bias from ignoring sampling variation in 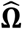 and 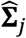 is negligibly small. A possible exception, not discussed so far, arises if MTAG is used for GWAS meta-analysis across a large number of cohorts (in which case *T* is large). MTAG may be valuable for that purpose because (i) it accounts for sample overlap and cryptic relatedness across cohorts and (ii) different cohorts often have phenotypic data from different measures, which may be imperfectly genetically correlated and have different heritabilities. For such applications, to reduce bias in the MTAG standard errors, we recommend imposing reasonable parameter restrictions on the 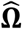 and 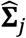 matrices (e.g., assuming that within groups of cohorts that analyzed identical phenotype measures, the heritability is equal and all pairwise genetic correlations are one).

#### 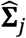 accurately captures sample overlap

MTAG relies on bivariate LD score regression (and by extension its assumptions) to estimate the correlation in GWAS estimation error due to sample overlap. To gauge MTAG’s performance, we simulate an extreme case of sample overlap using real data from the UK Biobank (UKB). We run three GWASs of height, each using two-thirds of the data, with 50% overlap between each pair of GWAS samples. Then we run MTAG on the three GWASs. Figure 2a is a scatterplot of the resulting MTAG *z*-statistics against the *z*-statistics from a single GWAS run on the entire UKB sample. Figure 2b is the scatterplot from an analogous analysis of DEP in UKB. The regression slope and *R*^2^ are both essentially one for both phenotypes, indicating that MTAG generates the correct *z-*statistics in these cases. **Supplementary Figure 2.1** shows that the results are similar when we repeat this analysis using four other phenotypes.

**Fig. 2.**
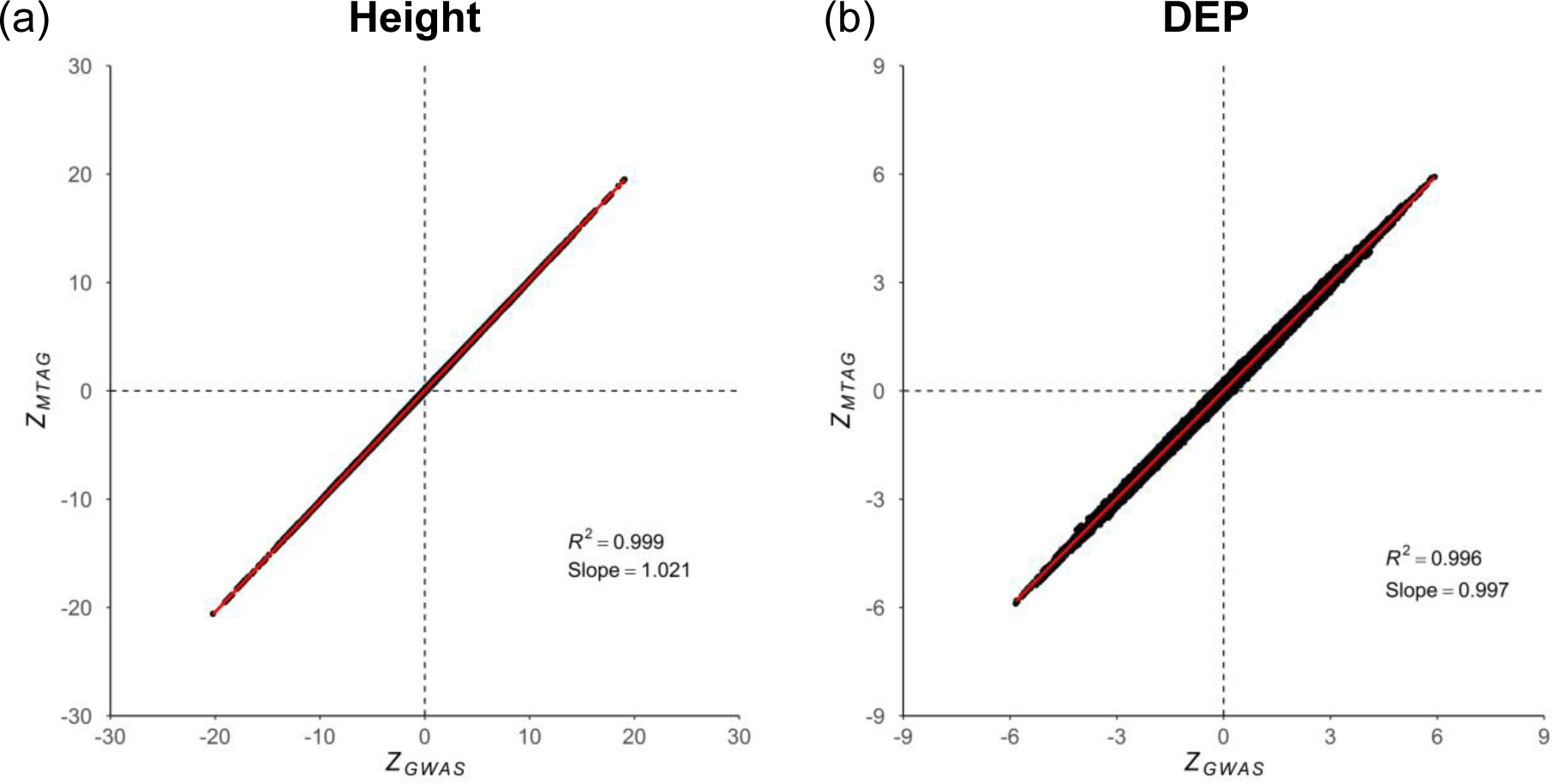
Evaluation of MTAG’s standard errors when there is sample overlap. The *x*-axis is a SNP’s *z*-statistic from a baseline GWAS conducted in UK Biobank. The *y*-axis is a SNP’s *z*-statistic from applying MTAG to three GWASs of each trait conducted on equally sized subsamples of the baseline sample, in which every pair of samples has 50% overlap. (**a**) Height. (**b**) Depressive symptoms. The figure illustrates near-perfect alignment. See **Supplementary Note**for details and results from analogous analyses on additional phenotypes.

### GWAS Summary Statistics for Depression, Neuroticism, and Subjective Well-Being

For our empirical application of MTAG, we build on a recent study by the Social Science Genetic Association Consortium (SSGAC) of three traits that have been found to be highly polygenic and strongly genetically related: depressive symptoms (DEP), neuroticism (NEUR), and subjective well-being (SWB). In these analyses, we combine data from the largest previously published studies^7–9,11^ with new genome-wide analyses from the genetic testing company 23andMe, Inc., and the first release of the UK Biobank (UKB) data. Relative to the previous SSGAC study, we reran the association analyses in UKB using a slightly revised analysis protocol, and much more importantly, we expanded the SSGAC meta-analyses for DEP and SWB. For DEP, we added the results from a recently published GWAS of depression in a large 23andMe cohort^7^, and for SWB, we added new association analyses of SWB in a 23andMe cohort. As depicted in Figure 3, there is substantial overlap between the estimation samples for the three traits. For additional details, see **Online Methods** and **Supplementary Note**.

**Fig. 3.**
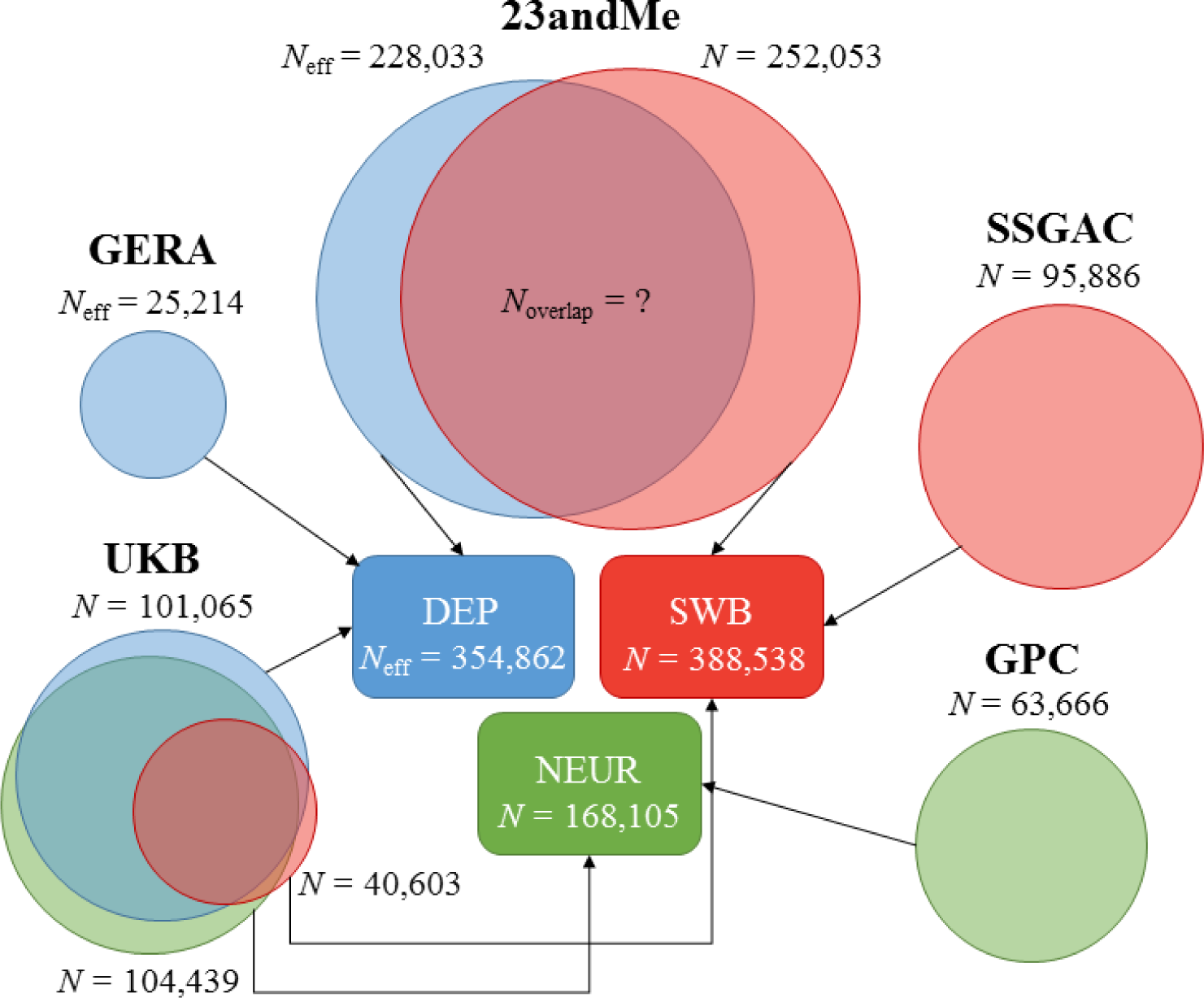
Cohorts included in GWAS meta-analyses for DEP, NEUR, and SWB. In UKB, the sample overlap in the summary statistics across the traits is known, whereas in 23andMe, the sample overlap in the summary statistics is unknown. MTAG accounts for both sources of overlap. SSGAC results,^20^ GPC results,^19^ GERA results,^18^ and 23andMe results for DEP^21^ all come from previously published work. The data from 23andMe for SWB are newly analyzed data for this paper. Data from the UKB for all three traits has been previously published,^20^ although we re-analyze it in this paper with slightly different protocols. *N*_eff_ is used instead of *N* when the cohort has case-control data (**Supplementary Note**). The sample size listed for each cohort corresponds to the maximum sample size across all SNPs available for that cohort. The total sample size for each trait corresponds to the maximum sample size among the SNPs available after applying MTAG filters. For details, see **Supplementary Note**.

### MTAG Results

We applied MTAG to the summary statistics from the three single-trait analyses described above. To enable a fair comparison between the MTAG and GWAS results, we restrict all analyses to a common set of SNPs (see **Online Methods** for details and recommended filters for MTAG).

Figure 4 shows side-by-side Manhattan plots from the GWAS and MTAG analyses for each trait. Approximately independent genome-wide significant SNPs, hereafter “lead SNPs,” were defined by clumping with an *R*^2^ threshold of 0.1 (**Online Methods**). From GWAS to MTAG, the number of lead SNPs increases from 32 to 64 for DEP, from 9 to 37 for NEUR, and from 13 to 49 for SWB. A list of all clumped SNPs with a *P* value less than 10^−5^ is in **Supplementary Table 4.5**.

**Fig. 4.**
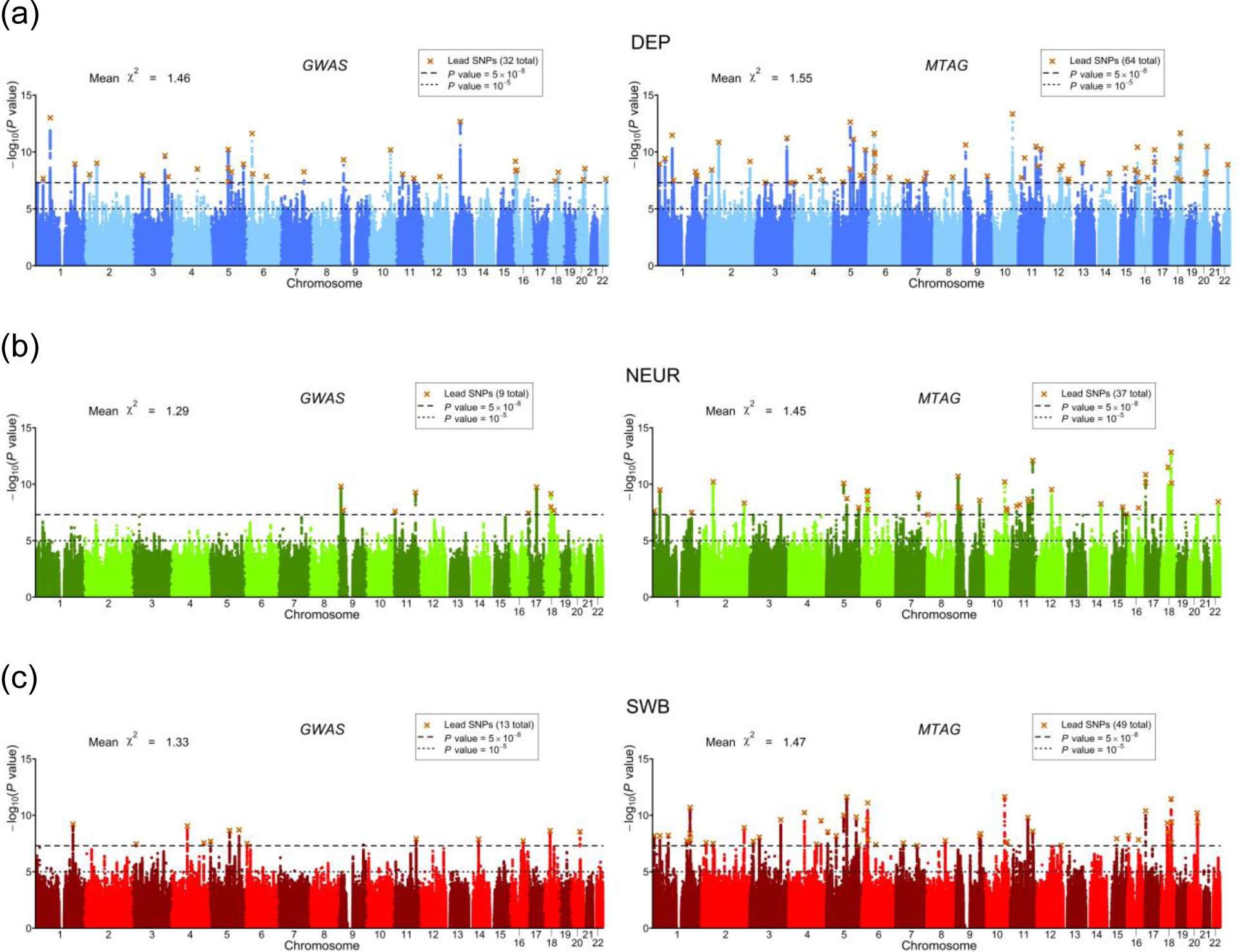
Manhattan plots of GWAS and MTAG results. (**a**) DEP, (**b**) NEUR, (**c**) SWB. The left and right plots display the GWAS and MTAG results, respectively, for a fixed set of SNPs. The *x*-axis is chromosomal position, and the *y*-axis is the significance on a −log_10_ scale. The upper dashed line marks the threshold for genome-wide significance (*P* = 5 × 10^−8^), and the lower line marks the threshold for nominal significance (*P* = 10^−5^). Each approximately independent genome-wide significant association (“lead SNP”) is marked by ×. The mean *χ*^2^-statistic across all SNPs included in the analysis is displayed in the top left corner of each plot.

For the MTAG hits, we calculate the maxFDR assuming that at least 10% of SNPs are non-null for each trait (our estimates of the actual percentage non-null are 59-65% across the three traits; see **Online Methods**). The maxFDR is 0.0014 for DEP, 0.0080 for NEUR, and 0.0044 for SWB. This calculation suggests that the hits are unlikely to be an artifact of the homogeneous-**Ω** assumption.

For each trait, we assess the gain in average power from MTAG relative to the GWAS results by the increase in the mean *χ*^2^-statistic. We use this increase to calculate how much larger the GWAS sample size would have to be to attain an equivalent increase in expected *χ*^2^ (**Online Methods**). We find that the MTAG analysis of DEP, NEUR, and SWB yielded gains equivalent to augmenting the original samples sizes by 27%, 55%, and 55%. The resulting GWAS-equivalent sample sizes are thus 449,649 for DEP, 260,897 for NEUR, and 600,834 for SWB.

### Replication of MTAG-identified Loci

To test the lead SNPs for replication, we use the Health and Retirement Study (HRS) and the National Longitudinal Study of Adolescent to Adult Health (Add Health), which both contain high-quality measures of DEP, NEUR, and SWB. Because HRS was included in the SSGAC discovery sample for SWB, we re-ran the GWAS and MTAG analyses for SWB after omitting it. Although our replication samples are too small for well-powered replication analyses of single-SNP associations, we are well powered to test the SNPs jointly. For the set of MTAG-identified lead SNPs for each trait, we regressed the effect sizes in HRS and in Add Health on the MTAG effect sizes, after correcting the MTAG effect-size estimates for the winner’s curse (**Online Methods**). The regression slope for each replication cohort was then meta-analyzed. If the SNP effect sizes taken altogether replicate, then we expect a slope of one. The regression slopes are 0.88 (s.e. = 0.22) for DEP, 0.76 (s.e. = 0.21) for NEUR, and 0.99 (s.e. = 0.33) for SWB (Figure 5). In all cases, the slope is statistically significantly greater than zero (one-sided *P* = 2.16 × 10^−5^, 1.87 × 10^−4^, and 1.52 × 10^−3^, respectively) but not statistically distinguishable from one.

**Fig. 5.**
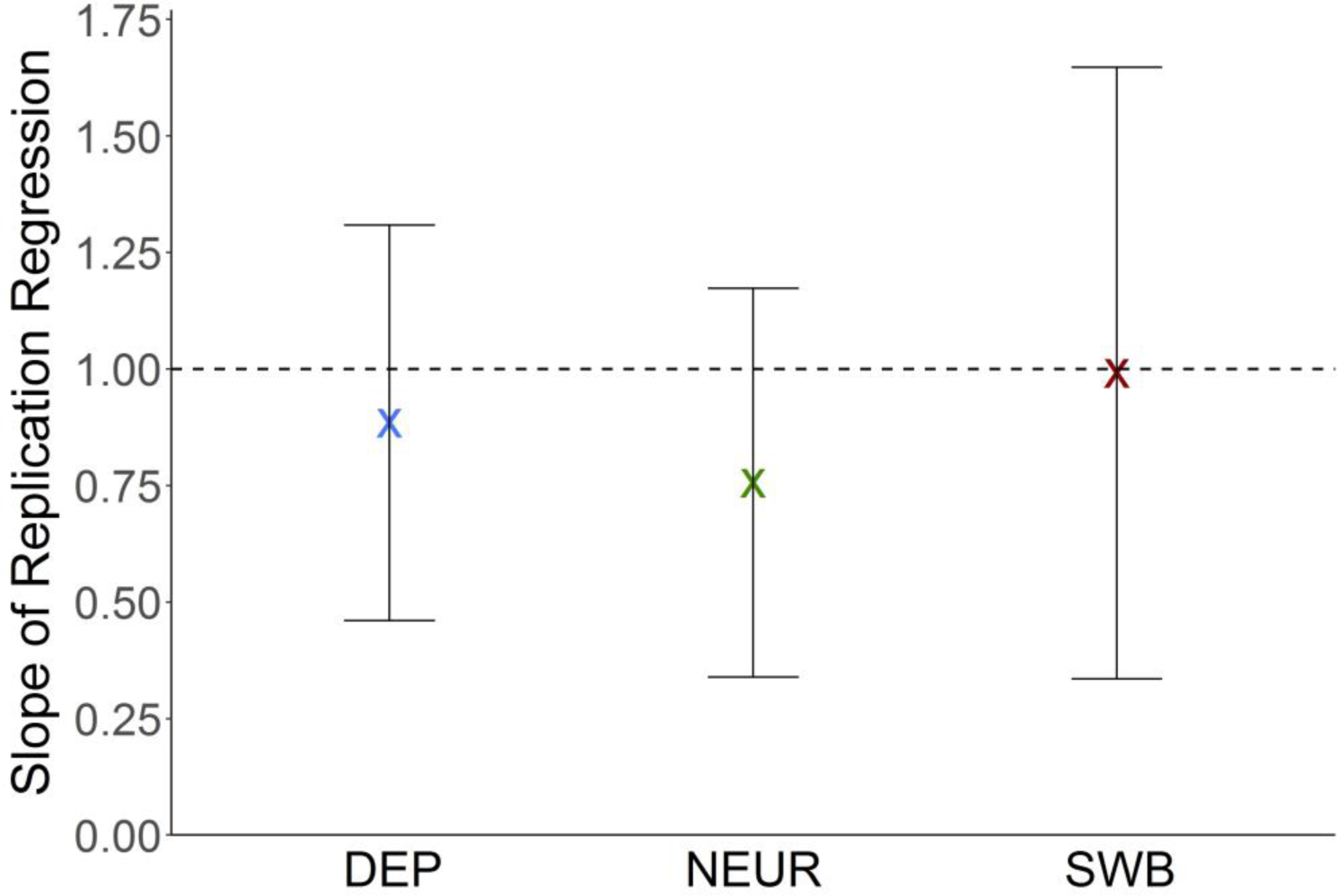
Regression-based test of replicability of MTAG-identified loci. For each trait and in each of two independent replication cohorts (HRS and Add Health, combined *N* = 12,641), we regressed the estimated effect sizes of the MTAG-identified loci on their winner’s-curse-adjusted MTAG effect sizes. The intercept is constrained to zero in these regressions. The plotted regression coefficients are the sample-size-weighted means across the replication cohorts, with 95% intervals. See **Supplementary Note** for details and cohort-level results.

### Polygenic Prediction

We next compare the predictive power of polygenic scores constructed from GWAS versus MTAG association statistics. We again use the HRS and Add Health as our prediction samples (and we obtain the SNP effect estimates for SWB from the analyses that omit HRS from the discovery sample).

We measure the predictive power of each polygenic score by its incremental *R*^2^, defined as the increase in coefficient of determination (*R*^2^) as we move from a regression of the trait only on a set of controls (year of birth, year of birth squared, sex, their interactions, and 10 principal components of the genetic data) to a regression that additionally includes the polygenic score as an independent variable.

Figure 6 and Table 1 summarize the results from our pooled analysis of Add Health and HRS. The GWAS-based polygenic scores have incremental *R*^2^’s of 1.00% for DEP, 1.27% for NEUR, and 1.20% for SWB. The corresponding MTAG-based polygenic scores all have greater predictive power: 1.17% for DEP, 1.65% for NEUR, and 1.57% for SWB. The proportional improvement in incremental *R*^2^ is in the range 17-30% for each trait, with 95% confidence intervals that do not overlap zero. The absolute levels of predictive power are clearly too small to be of clinical utility, but the improvements in *R*^2^ are close to those we would expect theoretically based on the observed increases in mean *χ*^2^-statistics (see **Online Methods**). Polygenic scores based on trait-specific MTAG results have greater predictive power than scores based on MTAG results for the other traits (Figures 6c and 6d), consistent with the theoretical result that MTAG results can be interpreted as trait-specific estimates.

**Table 1.** Summary of comparative analyses of GWAS and MTAG results

|  | DEP |  | NEUR |  | SWB |  |
| --- | --- | --- | --- | --- | --- | --- |
|  | GWAS | MTAG | GWAS | MTAG | GWAS | MTAG |
| <u>SNP-based comparisons</u> |  |  |  |  |  |  |
| Lead SNPs<br>($P < 5 \times 10^{-8}$) | 32 | 64 | 9 | 37 | 13 | 49 |
| Mean $\chi^2$ | 1.43 | 1.55 | 1.29 | 1.45 | 1.30 | 1.47 |
| $N_{\text{eff}}$ | 354,861 | 449,649 | 168,105 | 260,897 | 388,538 | 600,834 |
| Polygenic score<br>incremental $R^2$ | 1.00% | 1.17% | 1.27% | 1.65% | 1.20% | 1.57% |
| <u>Biological Annotation (DEPICT FDR < 0.05)</u> |  |  |  |  |  |  |
| # Prioritized Genes | 3 | 72 | 0 | 51 | 0 | 0 |
| # Gene Sets | 0 | 347 | 0 | 1 | 0 | 7 |
| # Tissues and cell<br>types | 10 | 22 | 0 | 21 | 0 | 12 |

**Fig. 6.**
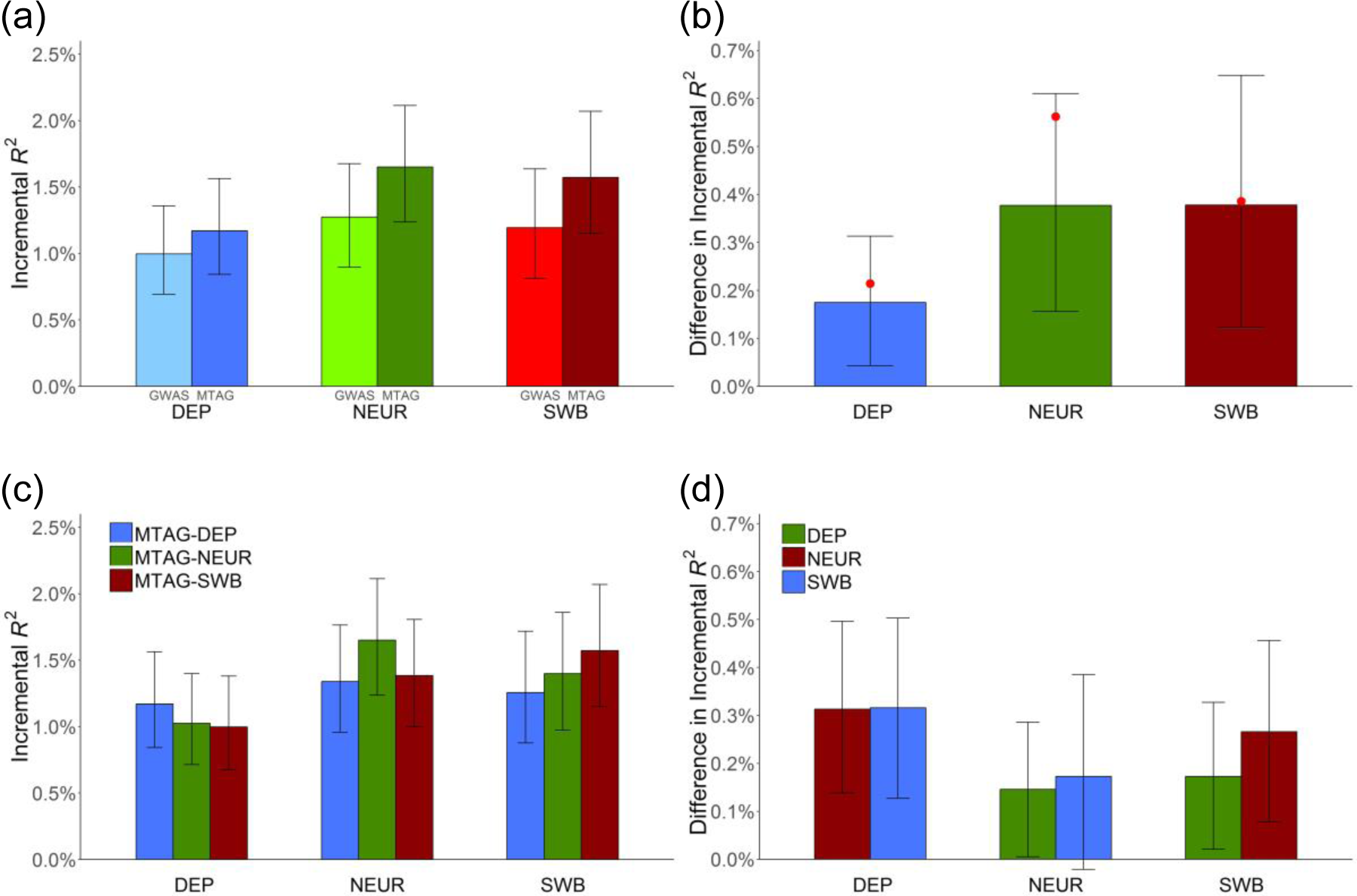
Predictive power of GWAS- and MTAG-based polygenic scores. Incremental *R*^2^ is the increase in *R*^2^ from a linear regression of the trait on the polygenic score and covariates, relative a linear regression of the trait on only covariates. The plotted incremental *R*^2^’s (and differences in incremental *R*^2^’s) are the sample-size-weighted means across the replication cohorts (HRS and Add Health, combined *N* = 12,641), with 95% intervals. See **Supplementary Note** for details and cohort-level results. (**a**) Incremental *R*^2^ of MTAG-based and GWAS-based polygenic scores. (**b**) Difference in incremental *R*^2^ between the GWAS- and the MTAG-based PGS. Red dots indicate the theoretically predicted gains in prediction accuracy (**Online Methods**). (**c**) Incremental *R*^2^ of polygenic scores constructed from the MTAG results for the predicted trait (“own-trait score”) or MTAG results for each of the other traits (“other-trait score”). The *x*-axis indicates the trait being predicted, and the bar color indicates which trait’s polygenic score is used. (**d**) Difference in incremental *R*^2^ between own-trait scores and the mean of the incremental *R*^2^’s from the other-trait scores.

### Biological Annotation

For a final comparison, we analyze both the GWAS and MTAG results using the bioinformatics tool DEPICT^15^. We present the prioritized genes, enriched gene sets, and enriched tissues identified by DEPICT at the standard FDR threshold of 5%.

Table 1 summarizes the results (see **Supplemental Tables 7.1** to **7.10** for the complete set of findings). In the GWAS-based analysis, very little enrichment is apparent. For DEP, 3 genes are identified, but no gene sets and only 10 tissues. For NEUR and SWB, no genes, gene sets, or tissues are identified. In contrast, the MTAG-based analysis is more informative. The strongest results are again for DEP, now with 72 genes, 347 gene sets, and 22 tissues. For NEUR, there are 51 genes, 1 gene set, and 21 tissues, and for SWB, zero genes, 7 gene sets, and 12 tissues.

For brevity, we discuss the specific results only for DEP; the results for NEUR and SWB are similar but more limited. For the tissues tested by DEPICT, Figure 7a plots the *P* values based on both the GWAS and MTAG results. As expected, nearly all of the enrichment of signal is found in the nervous system. To facilitate interpretation of the enriched gene sets, we used a standard procedure^16^ to group the 347 gene sets into ‘clusters’ defined by degree of gene overlap. Many of the resulting 46 clusters, shown in Figure 7b, implicate communication between neurons (‘synapse,’ ‘synapse assembly,’ ‘regulation of synaptic transmission,’ ‘regulation of postsynaptic membrane potential’). This evidence is consistent with that from the DEPICT-prioritized genes, many of which encode proteins that are involved in synaptic communication. For example, *PCLO*, *BSN*, *SNAP25*, and *CACNA1E* all encode important parts of the machinery that releases neurotransmitter from the signaling neuron.^17^

**Fig. 7.**
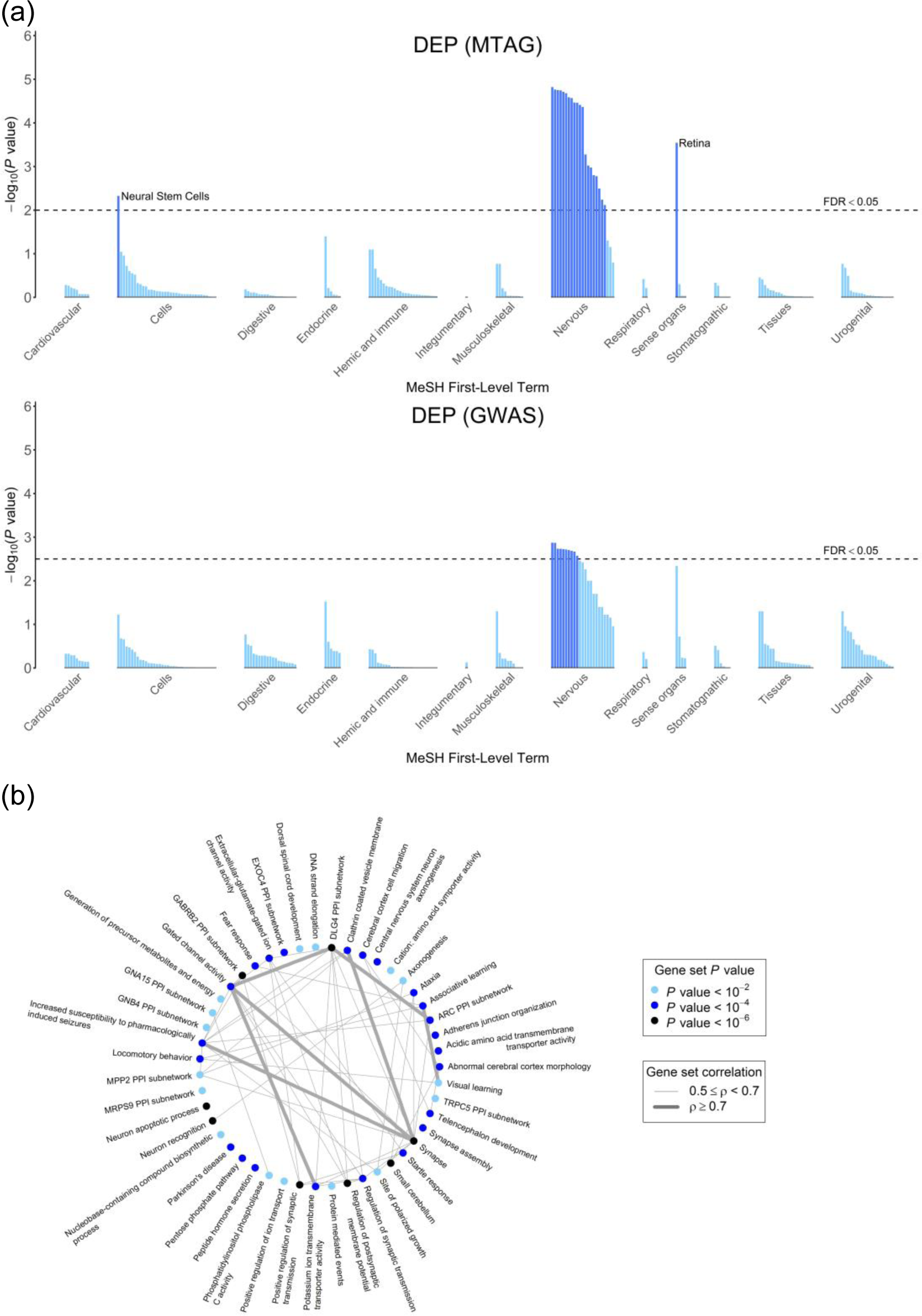
Biological annotation for DEP using the bioinformatics tool DEPICT. Results of the tissue-enrichment analysis based on the GWAS and MTAG results. The *x*-axis lists the tissues tested for enrichment, grouped by the location of the tissue. The *y*-axis is statistical significance on a −log_10_ scale. The horizontal dashed line corresponds to a false discovery rate of 0.05, which is the threshold used to identify prioritized tissues. (b) Gene-set clusters as defined by the Affinity Propagation algorithm23 over the gene sets from the MTAG results. The algorithm names clusters after an exemplary member of the gene set. The color of the point signifies the *P* value of the most significant gene set in the cluster. The line thickness between the gene-set clusters corresponds to the correlation between the named gene sets for each pair of clusters.

The results contain some intriguing findings. For example, while hypotheses regarding major depression and related traits have tended to focus on monoamine neurotransmitters, our results as a whole point much more strongly to glutamatergic neurotransmission. Moreover, the particular glutamate-receptor genes prioritized by DEPICT (*GRIK3*, *GRM1*, *GRM5*, and *GRM8*) suggest the importance of processes involving communication between neurons on an intermediate timescale,^18,19^ such as learning and memory. Such processes are also implicated by many of the enriched gene sets, which relate to altered reactions to stress and novelty in mice (e.g., ‘decreased exploration in a new environment,’ ‘increased anxiety-related response,’ ‘behavioral fear response’).

### Comparison to Other Multi-Trait Methods

We compared MTAG to three multi-trait methods that can be applied to an arbitrary number of GWAS summary with unknown overlap^12,13^ (see **Online Methods** and **Supplementary Note**). Unlike MTAG, these methods do not provide trait-specific SNP effect estimates but instead test whether the SNP is associated with none of the traits. We generate a (conservative) MTAG-based test of the same null hypothesis by using the minimum of the trait-specific MTAG *P* values, Bonferroni-adjusted for the number of traits. In two-trait simulations, we find that MTAG has greater power when the correlation in true effect sizes or GWAS estimation error is non-zero, especially when the traits’ GWASs are higher powered. In real-data applications to (i) DEP, NEUR, and SWB, and (ii) six anthropometric traits, MTAG identifies more loci. We test the anthropometric loci in GIANT consortium results and find that the loci identified by MTAG and missed by the other methods replicated at a higher rate than the loci identified by one of the other methods and missed by MTAG.

## DISCUSSION

We have introduced MTAG, a method for conducting meta-analysis of GWAS summary statistics for different traits which is robust to sample overlap. Both our theoretical and empirical results confirm that MTAG can increase the statistical power to identify *trait-specific* genetic associations. In our empirical application to DEP, NEUR, and SWB, we found that relative to the separate GWASs for the traits, MTAG led to substantial improvements in number of loci identified, predictive power of polygenic scores, and informativeness of a bioinformatics analysis. Table 1 summarizes the gains from MTAG across these analyses.

Because large-scale GWAS summary statistics are accessible for an ever-increasing number of traits and tools are now available for using summary statistics to easily identify clusters of genetically correlated traits,^20^ there will be many sets of traits to which MTAG could be applied. Which potential applications will be most fruitful? Our theoretical results indicate that, relative to the single-trait GWASs, MTAG will improve polygenic prediction quite generally. For identifying individual loci, MTAG will yield the greatest gains in statistical power and little inflation of the FDR for traits with high genetic correlation. We caution, however, that the FDR can become substantial if MTAG is applied to a large number of low-powered GWASs or to GWASs that differ a great deal in power—conditions that do not apply to our empirical application here. In all applications of MTAG, we recommend conducting FDR calculations and, of course, conducting replication analyses if possible.

## URLs

Social Science Genetic Association Consortium (SSGAC) website: http://www.thessgac.org/#!data/kuzq8.

MTAG software available at: https://github.com/omeed-maghzian/mtag.

## ACKNOWLEDGMENTS

We thank Jonathan Beauchamp, Philipp Koellinger, Örjan Sandewall, Carl Shulman, and Ronald de Vlaming for helpful comments and Rebecca Royer for research assistance. This research was carried out under the auspices of the Social Science Genetic Association Consortium (SSGAC). The study was supported by the Ragnar Söderberg Foundation (E9/11 E42/15), the Swedish Research Council (421-2013-1061), The Jan Wallander and Tom Hedelius Foundation, an ERC Consolidator Grant (647648 EdGe), the Pershing Square Fund of the Foundations of Human Behavior, the National Science Foundation’s Graduate Research Fellowship Program (DGE 1144083), and the NIA/NIH through grants P01-AG005842, P01-AG005842-20S2, P30-AG012810, and T32-AG000186-23 to NBER and R01-AG042568-02 to the University of Southern California. This research has also been conducted using the UK Biobank Resource under Application Number 11425. We thank the research participants and employees of 23andMe for making this work possible. We also thank Kathleen Mullan Harris and Add Health for early access to the data used in our replication and prediction analyses. A full list of acknowledgments is provided in the **Supplementary Note**.

## CONTRIBUTOR LIST FOR THE 23andMe RESEARCH TEAM

Michelle Agee, Babak Alipanahi, Adam Auton, Robert K. Bell, Katarzyna Bryc, Sarah L. Elson, Pierre Fontanillas, Nicholas A. Furlotte, David A. Hinds, Bethann S. Hromatka, Karen E. Huber, Aaron Kleinman, Nadia K. Litterman, Matthew H. McIntyre, Joanna L. Mountain, Carrie A.M. Northover, J. Fah Sathirapongsasuti, Olga V. Sazonova, Janie F. Shelton, Suyash Shringarpure, Chao Tian, Joyce Y. Tung, Vladimir Vacic, Catherine H. Wilson, and Steven J. Pitts.

## AUTHOR CONTRIBUTIONS

B.M.N., D.J.B., D.C. and P.T. oversaw the study. The theory underlying MTAG was conceived of and developed by P.T., with contributions from B.M.N., D.J.B., D.C., D.L., O.M., P.M.V. and R.K.W. O.M., P.T., and R.K.W. performed the simulations and developed the MTAG software. P.T. and P.M.V. designed the analyses comparing the observed MTAG gains to theoretical expectations. A.O., M.Z., R.W., O.M., and T.N. played major roles in data analyses. J.J.L. designed and executed the bioinformatics analyses. D.J.B., D.C., and P.T. coordinated the writing of the manuscript. All authors provided input and revisions for the final manuscript.

## CORRESPONDING AUTHORS

P.T., B.M.N., D.C., or D.J.B.

## ONLINE METHODS

This article is accompanied by a **Supplementary Note** with further details.

### Theory

There are *T* traits, which may be binary or quantitative. We standardize each trait and the genotype for each single-nucleotide polymorphism (SNP) *j* so that they all have mean zero and variance one. The length-*T* vector of marginal (i.e., not controlling for other SNPs), true effects of SNP *j* on each of the traits is denoted ***β**_j_*. We assume that these are random effects with mean **0** and variance-covariance matrix **Ω** that is the same across *j*. The mean is zero because we treat the choice of reference allele as arbitrary. We make the common assumption^14,21,22^ that the ***β**_j_*’s are identically distributed across *j*. The assumption implies that the expected amount of phenotypic variance explained is equal for each SNP, regardless of SNP characteristics such as allele frequency.

The length-*T* vector of GWAS estimates is denoted 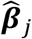, which is equal to the true effect vector plus estimation error, ***β**_j_* + ***ε**_j_*. The estimation error is the sum of sampling variation and biases (such as population stratification or technical artifacts). With any standard GWAS estimator (such as OLS or logistic regression), sampling variation is uncorrelated with ***β**_j_*. We assume that the biases are also uncorrelated with ***β**_j_*. The variance-covariance matrix of ***ε**_j_*, denoted **Σ**_*j*_, may differ across SNPs *j* due to differences in the SNPs’ sample sizes per trait and the SNPs’ sample overlap between traits, although we only account for the former in our estimation of **Σ**_*j*_.

MTAG is a generalized method of moments (GMM) estimator. To obtain the key moment conditions we will use, we consider the best linear prediction of the GWAS estimate for trait 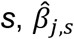, from the SNP’s true effect on trait *t*, *β_j,t_*. We use a first-order condition of this best linear prediction as the moment condition for trait 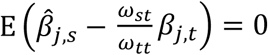, where *ω_st_* is the (*s, t*)^th^ element of **Ω**. There are *T* such moment conditions for *s* = 1,2, …, *T*, giving us the vector of moment conditions:

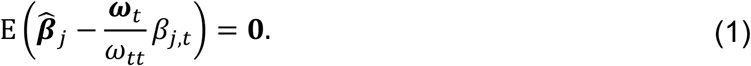

where ***ω**_t_* is a vector equal to the *t*^th^ column of **Ω**. Although *β_j,t_* is a random effect, we aim to estimate its (unknown) realized value. The efficient GMM estimator for *β_j,t_* based on the vector of moment conditions in equation (1) solves

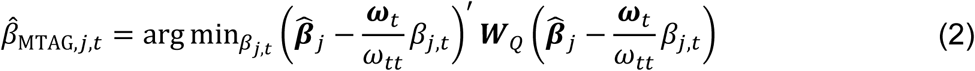

where 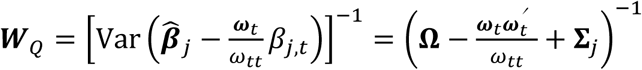 is the efficient weight matrix. Intuitively, the GMM estimator chooses the value of *β_j,t_* that minimizes a weighted sum of the squared deviations from the moment conditions, with deviations weighted more heavily if they are estimated more precisely. In the **Supplementary Note**, we show that the solution to the minimization problem in equation (2) is:

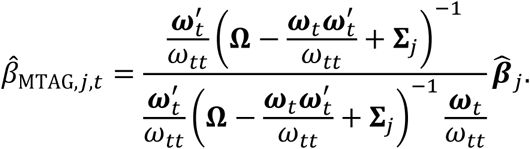

Standard asymptotic properties of GMM relate to *T* → ∞. In the **Supplementary Note**, we show that for fixed number of traits *T*, as the sample size for the GWAS of any trait *t* becomes large, the MTAG estimator 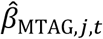 is consistent and asymptotically normal.

The sampling variance of the estimator is

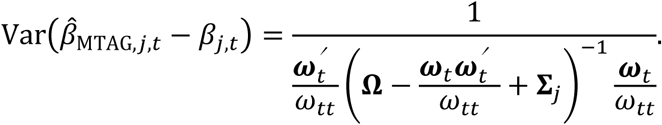

For each trait *t*, the standard error of the estimate is the square root of this quantity. As is standard, we obtain a *P* value using the fact that in large samples, 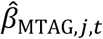 divided by its standard error follows a standard normal distribution under the null hypothesis.

Because of the homogeneous-**Ω** assumption, the above formulas for the MTAG estimator and its standard error effectively use the variance-covariance matrix of true SNP effects across all SNPs, **Ω**, to calculate the MTAG estimate for each SNP. If in fact there are different types of SNPs characterized by different variance-covariance matrices, then the MTAG estimator remains consistent but could be made more efficient if it took into account the different types of SNPs. In addition, the standard error formula is conservative on average across SNPs, which reduces MTAG’s statistical power to identify truly associated SNPs. Most importantly, the MTAG estimator is in general biased in finite samples, and it is biased away from zero for SNPs that are truly null, which causes the false positive rate to be inflated.

For each SNP *j*, given Σ_*j*_, the matrix **Ω** is estimated using the method of moments (see the **Supplementary Note** for discussion of the relationship to GMM). For each (*t, s*)^th^ entry of **Ω**, *Ω_ts_*, we use the moment condition 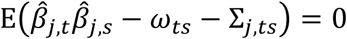. This moment condition is derived from observing that 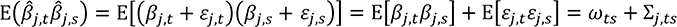. The estimator simply replaces the population expectation with the sample mean:

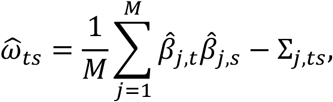

where *M* is the number of SNPs in the analysis. Intuitively, the estimated covariance in true genetic effects between trait *t* and trait *s* is equal to the covariance in their observed GWAS coefficients minus the covariance in GWAS coefficients that is due to correlated estimation error.

For expositional simplicity, our derivations above and in **Supplementary Note** are parameterized in terms of the parameter vector 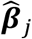. We note, however, that the input to the MTAG software is the standard output from meta-analysis software: *z-*statistics and sample sizes. Because MTAG is applied to *z*-statistics, the GWAS summary statistics do not need to have been estimated using traits and genotypes that were standardized.

### Special Cases

There are three special cases of MTAG that may often be relevant in practice and for which the estimation procedure is made faster and more efficient. The MTAG software offers the option to specialize the analysis for these cases.

#### No sample overlap across traits

In this case, the off-diagonal elements of Σ_*j*_ can be set equal to zero, so LD score regression needs to be run only *T* rather than *T*(*T* + 1)/2 times. Note that this version of MTAG does not take into account correlation in estimation error across traits that is due to bias. For this reason, LD score regression should be run on the MTAG results, and the resulting MTAG standard errors should be inflated by the square root of the estimated intercept.

#### Perfect genetic correlation but different heritabilities

This case arises when the “traits” are different measures of the same trait, some with more measurement error than others, or when the variance in the trait due to non-genetic factors differs. Here the **Ω** matrix has only *T* rather than *T*(*T* + 1)/2 unique parameters to be estimated.

#### Perfect genetic correlation and equal heritabilities

This special case corresponds to the “traits” being (the same measure of) a single trait; in other words, applying MTAG instead of inverse-variance-weighted meta-analysis to *T* GWAS results. Doing so can be useful if there is sample overlap in the GWAS results. In this case, as noted in the main text, MTAG specializes to 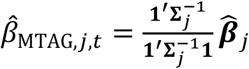 for all *t*, and it is no longer necessary to estimate **Ω**.

### Computational Run-time

MTAG is computationally quick because all its steps have closed-form solutions. In the real-data application described in this paper—with three traits and 6.1M SNPs—the median run time across five identical runs using one core of a 2.20 GHz Intel(R) Xeon(R) CPU E5-2650 v4 processor was approximately 28 minutes. Of course, run time may vary as a function of the computing environment.

### MTAG’s Genome-Wide Mean Squared Error (MSE)

The genome-wide MSE of the MTAG estimates is simply equal to their sampling variance (given above):

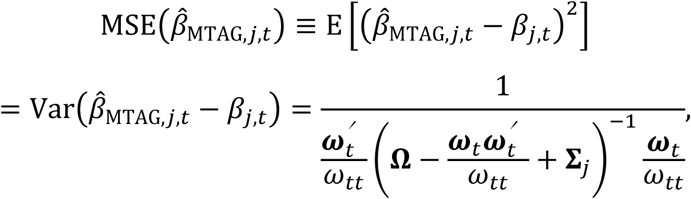

where the first equality follows because both the true effects *β_j,t_* and the MTAG estimates 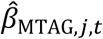 are mean zero. This formula for the MSE holds very generally; in particular, it does not require assuming that **Ω** is homogeneous across SNPs (because the genome-wide MSE is a property regarding the mean across all the SNPs included in the analysis). In the formula, **Ω** is (re-)defined as the genome-wide (i.e., across-SNP) variance-covariance matrix of the SNPs’ true effects on the traits. In **Supplementary Note**, we show that the MSE of the MTAG estimates are always weakly smaller than the MSE of the corresponding single-trait GWAS estimates, which equals 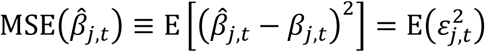.

### MTAG’s Power and False Discovery Rate (FDR) When Effect Sizes Are Mixture-Normal Distributed

Suppose that the vector of SNP *j*’s effects on the traits *β_j_* is drawn from a mixture of mean-zero multivariate normal distributions. The distribution of component *c* = 1,2, …, *c* is *β_j_*|*c* ∼ *N*(**0, Ω**_*c*_), and its mixture weight is denoted *p_c_*, where 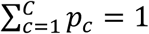. In this case, the *z*-statistic associated with the MTAG estimate 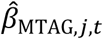 is a mixture distribution with component distributions

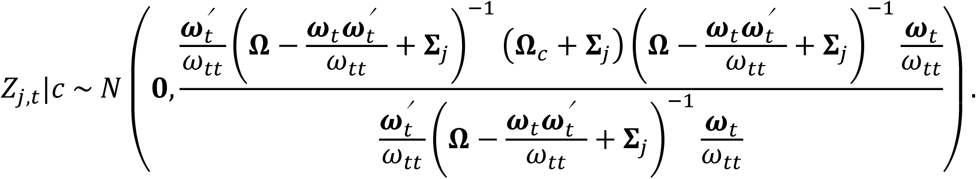

To define power and FDR, let *D* denote the set of components such that a SNP is null for trait *t* (i.e., the *t*^th^ element of *β_j_* is drawn from a degenerate distribution with all mass on 0). Power for trait *t* can be calculated as

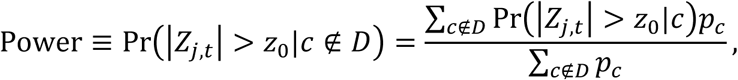

where *z*_0_ is the *z*-statistic associated with genome-wide significance. The FDR for trait *t* can be calculated as

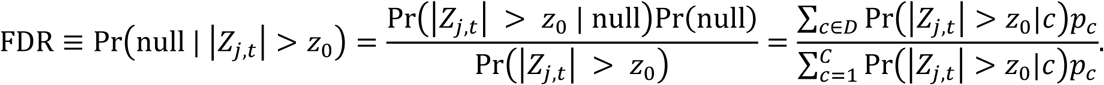

### Maximum FDR (MaxFDR) When Effect Sizes Are Multivariate Spike-and-Slab Distributed

Starting with the mixture-normal setup in the derivation of power and the FDR, we assume that there are *C* = 2^*T*^ components, corresponding to all possible combinations of the SNP being null for some subset of traits and non-null for the others. Let 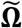 denote the variance-covariance matrix of true effect sizes for the component in which the SNP is non-null for all the traits. We assume that the variance-covariance matrix of true effect sizes for any component *c*, denoted **Ω**_*c*_, is equal to 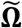 but with the rows and columns zeroed out that correspond to null traits in component *c*. Given our estimate of **Ω**, for any vector of mixing weights ***p*** = (*p*_1_, *p*_2_, …, *p_c_*), we construct an estimate of 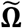: we set the (*t, s*)^th^ entry of 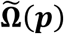 equal to 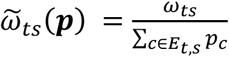, where *E_t,s_* is the set of components in which the SNP is non-null for both traits *t* and *s*. We call the mixing weights ***p*** *feasible* if the resulting matrix 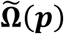 is positive semi-definite. We maximize the FDR (given by the formula above) over all feasible mixing weights ***p***. Given that the FDR may not be a unimodal function of ***p***, we maximize using a grid search. Since ***p*** has 2^*T*^ elements, it may be computationally infeasible to perform a fine grid search when *T* is larger than three or four traits.

### Simulations

To speed computations, instead of simulating data and then estimating effect sizes, we directly generated effect-size estimates by adding multivariate-normally-distributed noise to the simulated effect sizes. The variance of the noise for each trait was pinned down by the assumed GWAS expected *χ*^2^-statistics, and the covariance of the noise between the traits was pinned down by the assumed GWAS expected *χ*^2^-statistics and correlation of GWAS estimation error across traits.

In our simulations, we cannot estimate **Σ**_*j*_ using LD score regressions because we directly simulate effect sizes rather than data. Nonetheless, we would like to use a matrix for 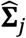 that contains the same amount of sampling variance that would have been present if we had simulated data and then ran LD score regressions. To accomplish this, in each replication we directly generated 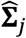 by adding noise to the true value of **Σ**_*j*_. The variance of the noise was calibrated against the LD-score-regression intercept standard errors for the GWAS results of DEP, NEUR, and SWB that we estimate in our empirical application but scaled to be larger or smaller when the simulated GWAS had more power (see **Supplementary Note**).

### GWAS Meta-analyses of DEP, NEUR, and SWB

Details on the cohorts, phenotype measures, genotyping, quality-control filters, and association models are provided in **Supplementary Note** and **Supplementary Table 3.1** to **3.4**. As shown in Figure 3, there is substantial overlap in samples across the three GWAS meta-analyses.

All analyses were based on autosomal SNPs from cohorts with genotypes imputed against the 1000 Genomes reference panel. The input files in each meta-analysis were subject to a uniform set of quality-control and diagnostic procedures. These are described in the previous SSGAC study^11^ and **Supplementary Note**.

As expected under polygenicity^23^, we observe inflation of the median test statistic in each GWAS (λ_GC,DEP_ = 1.36, λ_GC,NEUR_ = 1.24, λ_GC,SWB_ = 1.28; see **Supplementary Fig 3.2**). The intercept estimates from LD score regression are all below 1.02, however, suggesting that nearly all of the observed inflation is due to polygenic signal.^14^ When we report GWAS results, as in the SSGAC study^11^ we account for the potential bias due to this small amount of stratification by inflating the standard errors of our GWAS estimates by the square root of the LD score regression intercept.

Manhattan plots from each of the GWAS meta-analyses are shown in **Supplementary Figs 3.3a**, **b**, and **c**. Our NEUR meta-analysis was based on the same cohort-level data as the SSGAC study^11^ and unsurprisingly yielded substantively identical results: 10 lead SNPs. Consistent with what studies have reported for other complex traits, the increased discovery samples for DEP and SWB relative to the SSGAC study increased the number of lead SNPs: from 2 to 32 for DEP (*N*_eff_ = 149,707 to 354,862) and from 3 to 13 for SWB (*N* = 298,420 to 388,538). Applying bivariate LD score regression^6^ to the GWAS results, we estimate the genetic correlations to be 0.72 (s.e. = 0.026) between DEP and NEUR, -0.67 (s.e. = 0.027) between NEUR and SWB, and -0.69 (s.e. = 0.024) between DEP and SWB (see **Supplementary Table 3.6)**.

### Clumping Algorithm

We applied the same clumping algorithm to the GWAS and MTAG results to identify each set of “lead SNPs.” Our clumping algorithm is the same as in the previous SSGAC study.^11^ First, the SNP with the smallest *P* value was identified in the meta-analysis results. This SNP was designated the index SNP of clump 1. Second, we identified all SNPs on the same chromosome whose LD with the index SNP exceeds *R*^2^ = 0.1 and assigned them to clump 1. To generate the second clump, we removed the SNPs in clump 1 and then iterated the process to identify further index SNPs and their corresponding clumps until no SNPs remain.

### MTAG SNP Filters

Since the derivation of MTAG relies on some assumptions regarding features of the distributions of the effect sizes and estimation error, its performance may be sensitive to violations of those assumptions. To reduce the risk of extreme violations, when we apply MTAG, we impose three additional SNP filters beyond the standard filters used in a GWAS.

First, we restrict the set of SNPs to those with a minor allele frequency greater than 1%. This filter is motivated by the homogeneous-**Ω** assumption and by the assumption that each SNP explains the same amount of phenotypic variation in expectation. Rare variants may follow a different effect-size distribution both in terms of the variance and covariance of their effect sizes, which could bias the MTAG estimates.

Second, for each trait, we restrict variation in SNP sample sizes by calculating the 90^th^ percentile of the SNP sample-size distribution and removing SNPs with a sample size smaller than 75% of this value. This filter is similar to, though slightly more strict than, the sample-size filter recommended for LD Score regression.^14^ If a SNP’s effect is estimated in a relatively small subset of the sample, then the sample overlap across traits will likely be different for that SNP than for other SNPs in the sample. In that case, the covariance of the estimation error across traits as estimated by LD score regression may not be a good approximation to the covariance of the estimation error for that particular SNP.

Third, we drop SNPs in genomic regions containing SNPs that are outliers with respect to their effect-size estimates. Because the effect sizes of these SNPs appear to have a different variance-covariance matrix than the rest of the genome, including these regions would likely lead to the biases and inefficiencies that can occur when the homogeneous-**Ω** assumption is violated. In our empirical application, in the GWAS of NEUR, the effect sizes of SNPs in a region of chromosome 8 that tag an inversion polymorphism have been found to be strongly inflated relative to the effects estimated for SNPs in all other regions of the genome.^10,11^ Therefore, we omit SNPs in chromosome 8 between base-pair positions 7,962,590 and 11,962,591.

### GWAS-Equivalent Sample Size for MTAG

The increase in the mean *χ*^2^-statistic for each trait from the GWAS results to the MTAG results can be used to calculate a “GWAS-equivalent sample size” for MTAG. Under the assumptions of LD score regression,^14^ the expected *χ*^2^-statistic for some SNP with LD score *ℓ_j_* is

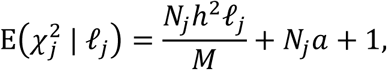

where *N_j_* is the sample size for SNP *j*; *h*^2^ is the SNP heritability of the trait; *M* is the number of SNPs for which we define the SNP heritability; and *a* is the variance due to biases (e.g., due to population stratification). Note that 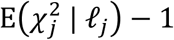 scales linearly with *N_j_* as long as *M* and *ℓ_j_* are held constant in the additional samples.^24–26^ Since the individuals included in all GWASs are of European ancestry, *M* and *ℓ_j_* are indeed expected to be approximately constant.^24–26^ Thus, we can use the mean *χ*^2-^ statistic from the GWAS and the MTAG results to calculate how much larger the GWAS sample size would have to be to give a mean *χ*^2^-statistic equal to that attained by MTAG:

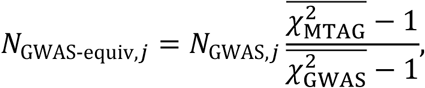

where 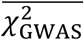 is the mean *χ*^2^-statistic in the GWAS results and 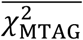 is the mean *χ*^2-^ statistic in the MTAG results. Put another way, conducting MTAG gives the same power (as measured by mean *χ*^2^-statistic) as conducting GWAS in sample size that is larger by a factor of 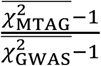.

For DEP, going from GWAS to MTAG, the mean *χ*^2^-statistic increases from 1.44 to 1.60, implying an increase in the GWAS-equivalent sample size by a factor of

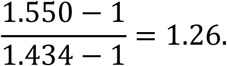

Thus, the MTAG analysis has statistical power equivalent to a GWAS of DEP conducted in 354,861 × 126% = 449,649 individuals. For NEUR, the mean *χ*^2-^statistic rises from 1.284 to 1.557, implying a GWAS-equivalent sample size for MTAG that is 96% larger than the GWAS sample size: the effective sample size is 168,105 × 196% = 329,835 individuals. For SWB, the mean *χ*^2^-statistics rises from 1.308 to 1.570, implying a GWAS-equivalent sample size 85% larger than the GWAS: 388,538 × 185% = 718,284 individuals.

### Expected Increase in Mean *χ*^2^-statistic from MTAG

The expected *χ*^2^-statistic of the GWAS summary statistics for trait *t* is

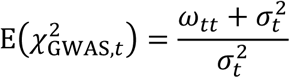

where 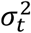 is the *t*^th^ diagonal element of **Σ**_*j*_. The expected *χ*^2^-statistic of the MTAG summary statistics for trait *t* is

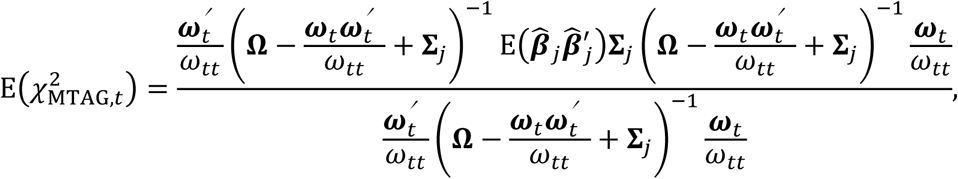

where we can substitute 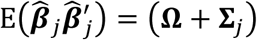.

Plugging our estimates 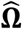 and 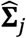into the above equations, we can use the expected *χ*^2^-statistics to calculate the theoretically expected gain in GWAS-equivalent sample size from applying MTAG (as derived previously):

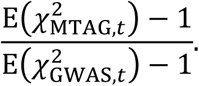

Note that 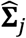 is a function of the sample sizes used to generate the GWAS summary statistics, and we use the 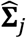 corresponding to the maximum sample size among the SNPs used in MTAG. Based on this formula, the theoretically expected increases in the GWAS-equivalent sample sizes are 28%, 58%, and 56% for DEP, NEUR, and SWB, respectively. These are very close to the observed increases of 27%, 55%, and 54%.

### Winner’s Curse Correction for Replication Analysis

MTAG estimates are corrected for winner’s curse following procedures previously described.^11^ Briefly, for each trait, we use maximum likelihood to fit the MTAG results to a (univariate) spike-and-slab distribution such that

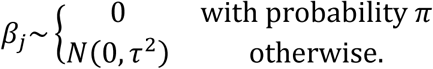

For DEP, NEUR, and SWB, we estimate 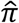 to be 0.598, 0.652, and 0.633 and 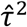 to be 3.12 × 10^−6^, 5.05 × 10^−6^, and 2.15 × 10^−6^, respectively. We then use these estimates as the parameters of the prior distribution and calculate the posterior distribution of the effect size *β_j_* given the estimate 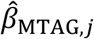 for each SNP as

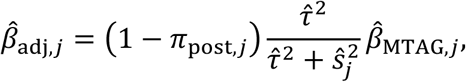

where *π_post,j_* is the posterior probability that *β_j_* = 0 and 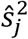 is the squared standard error of the MTAG estimate.

### Polygenic Prediction

We used the Health and Retirement Study^27^ (HRS) and the National Longitudinal Study of Adolescent to Adult Health (Add Health) as our prediction cohorts. We applied the same SNP filters as in the main MTAG analyses. Additionally, we restricted the set of SNPs used to construct the scores to HapMap3 SNPs for comparability across the two prediction cohorts. We calculated the SNP weights using the software package LDpred, assuming a fraction of causal SNPs equal to 1. The scores were constructed in PLINK using genotype probabilities obtained from 1000 Genomes imputation.

Bootstrapped confidence intervals were calculated by drawing, with replacement, a sample of equal size to the prediction sample, and then calculating the incremental *R*^2^ for the GWAS-based polygenic score, the MTAG-based polygenic score, and the difference between them. Our pooled results were obtained as a sample-size-weighted sum of HRS and Add Health results. As the bounds of the 95%-confidence intervals, we use the 2.5^th^- and 97.5^th^-percentile values of the incremental *R*^2^’s across 1000 bootstrap draws.

### Expected Increase in Polygenic-Score Predictive Power from MTAG

The phenotypic value of a trait in individual *i*, denoted *y_i_*, can be decomposed into the sum of the additive genetic variance component and a residual:

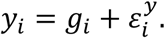

We denote the GWAS- and MTAG-based polygenic scores for the trait by 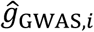 and 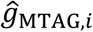, respectively. Note that GWAS and MTAG produce consistent estimates of the SNP effect sizes, and LDpred^22^ produces a consistent estimate of the additive genetic variance component. Therefore, each polygenic score *k* ∈ {GWAS, MTAG} is approximately equal to *g_i_* plus estimation error:

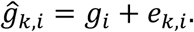

By the central limit theorem, the estimation error is approximately normally distributed,

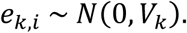

The variance *v_k_* is inversely proportional to the sample size as long as the effective number of chromosome segments, *M_e_*, is the same in every GWAS sample in the analysis.^24–26^ As in the calculation of the GWAS-equivalent sample size, where we assume that *M_e_* is the same in every GWAS sample and in the prediction sample, the expected predictive power of a polygenic score is

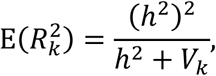

where *h*^2^ is the SNP heritability of the trait.^28–30^ (Note that if *M_e_* were to differ greatly across samples, then it would be important to take this into account when calculating the expected predictive power.^24,25^)

Using the GWAS results, we obtain an estimate of *h*^2^ using LD score regression^14^ and an estimate of 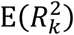 from the predictive power of the GWAS-based polygenic score. Plugging these estimates into the above formula, we solve for an estimate of *V*_GWAS_. We then multiply this value by 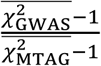 (which we showed previously is equal to the ratio of the GWAS sample size to the MTAG’s GWAS-equivalent sample size) to obtain an estimate of *V*_MTAG_. Substituting this back into the above formula along with our estimate of *h*^2^ gives us the expected predictive power of the MTAG-based PGS.

Results of this calculation are found in Panel C of **Supplementary Table 6.1**. For DEP, NEUR, and SWB, respectively, we anticipated increases in predictive power of 0.21, 0.56, and 0.39 percentage points. All three anticipated increases are within their respective estimated confidence intervals: [0.04, 0.31], [0.16, 0.61], and [0.12, 0.65]. Overall, the observed gains in predictive power relative to conventional GWAS-based polygenic scores are thus consistent with theoretical expectations.

### Biological Annotation

For the identification of genes, we supplemented the DEPICT inventory with protein-coding genes that have a status of ‘known’ in GENCODE (downloaded February 26, 2015). Specifically, we assigned such a gene to a lead SNP in **Supplementary Tables 7.1**, **7.4**, and **7.7** if it either encompasses the SNP in a DEPICT-defined locus or has the start site closest to such a SNP.

### Comparative Analyses

We conducted analyses comparing MTAG to other multi-trait methods that can be applied in the specific setting for which MTAG was developed: the only available inputs are summary statistics from an arbitrary number of genome-wide analyses conducted in samples with unknown overlap. We identified three methods satisfying these criteria: *S*_Hom_, *S*_Het_, and a method we call Bolormaa.^12,13^ Theoretically, we highlight two properties of MTAG that distinguish it. First, the alternative methods only provide an omnibus *P* value from the test of the joint null hypothesis that the SNP is not associated with any of the traits analyzed. In settings where the purpose of the multi-trait analysis is to test for association between a SNP and a *single* trait or to improve the prediction accuracy of a polygenic score, the alternative methods are not readily applicable. Second, the other methods lose efficiency by not distinguishing between variance and covariance in the GWAS test statistics that is due to true genetic signal versus estimation error. In the **Supplementary Note** we provide a more detailed theoretical overview and simulation evidence consistent with our theoretical conclusions (**Supplementary Fig. 8.1**).

We also performed comparative analyses based on summary statistics from actual GWASs. In our first such empirical application, we used summary statistics from the three single-trait GWASs (DEP, NEUR, and SWB) analyzed in this paper’s main MTAG analysis. In our second application, we applied MTAG and each of the three alternative methods to summary statistics from sex-stratified genome-wide analyses of height, body mass index (BMI) and waist-to-hip-ratio adjusted for BMI (WHRadjBMI) published by the GIANT consortium in 2013.^31^ In order to compare MTAG to the other methods, we use the Bonferroni *P* value of the joint null hypothesis: *P* = *T* × min{*P_t_*}, where *P_t_* is MTAG’s trait-*t P* value for the SNP. This *P* value is conservative because the trait-specific *P* values are in general correlated. Nonetheless, in both applications, we found that MTAG compares favorably in terms of the number of loci identified. We also leveraged more recent studies by the GIANT consortium to examine the replicability of the loci identified by the multi-trait methods.^32–34^ We found that MTAG-identified loci that evaded detection by the alternative methods appeared to replicate at a higher rate than loci identified by one of the alternative methods and missed by MTAG (**Supplementary Figs. 8.2-8.3** and **Supplementary Table 8.1**).

We did not compare MTAG to PleioPred,^4^ a new method that is closely related to MTAG, because it can be applied only to two traits and is designed only for polygenic prediction (not identification of SNP associations). We expect that PleioPred will outperform MTAG in polygenic prediction when the genetic architecture of the traits aligns with the spike-and-slab distribution assumed by the method, as seems to be the case for the empirical application studied in that paper.

